# *Hedyotis Diffusae Herba* Mitigates Rotenone-Induced Neurotoxicity via Akt Pathway in SH-SY5Y Cells

**DOI:** 10.1101/2025.08.03.668347

**Authors:** Jian Wang, Yisong Liu, Ying Fan, Meng Yuan, Wanying Xie, Xingchun Wang, Yi Zhong, Jiaxin Zhan, Xinyao Huang, Yibo Zhang, Junxi Long, Xingxia Wang

## Abstract

Hedyotis Diffusae Herba (HDH), known in traditional Chinese medicine for its heat-clearing and detoxifying properties, has been widely used to manage inflammation-related conditions. However, its neuroprotective potential and mechanism in the context of Parkinson’s disease (PD) remain unclear.Network pharmacology and molecular docking were applied to predict the interactions between HDH components and PD-associated targets. Molecular dynamics simulations assessed the binding stability of major compounds to core targets. Rotenone-induced SH-SY5Y cell injury served as an in vitro PD model to evaluate HDH’s biological effects. Cell viability, reactive oxygen species (ROS), IL-6 secretion, and Akt phosphorylation were assessed by CCK-8 assay, ROS detection kit, ELISA, and western blot, respectively. MK2206, an Akt inhibitor, was used to validate pathway involvement.Seven bioactive compounds were identified in HDH, among which stigmasterol and quercetin exhibited strong binding affinities with AKT1, IL-6, and JUN. Simulations confirmed stable interactions. HDH significantly increased cell viability and decreased intracellular ROS and IL-6 levels in rotenone-treated SH-SY5Y cells. Akt phosphorylation was restored by HDH, and the effect was blocked by MK2206, indicating Akt pathway participation.HDH exerts neuroprotective effects against rotenone-induced toxicity via antioxidant and anti-inflammatory activities and Akt pathway activation. These findings provide mechanistic insight and pharmacological support for HDH as a potential candidate in complementary therapy for PD.

## 1 Introduction

Parkinson’s disease (PD) ‘s incidence rate is between 1% and 2% and is the second most common neurodegenerative disease, with a global prevalence exceeding 6 million people. This figure is the result of a 2.5-fold increase in prevalence over the past 30 years, making PD one of the leading causes of neurological disability (Tolosa, Garrido, Scholz, & Poewe, 2021).The clinical hallmark of PD is a movement syndrome that is characterized by bradykinesia, resting tremors, and rigidity, in addition to changes in posture and gait. The movement disorders lead to progressive disability, impairment of activities of daily living, and a decline in the quality of life. At the same time, the development of postural instability, the increase in gait difficulties, as well as dysphagia and dysarthria drive the progression of the movement disorders (Jankovic, 2008).At the same time, mutations or variations in many genes can cause PD and increase the susceptibility to it. However, many of these aspects remain unresolved and there is a lack of conclusive research data(Pickrell & Youle, 2015).There is no cure for PD, and the best treatment options can only provide symptom relief and do not alleviate or reverse the progression of the disease. The quality of life, psychological burden, social situation, etc. of people with PD are all significantly affected scientists and relevant departments are in urgent need of finding new drugs to radically cure PD. Therefore, there is an urgent need for novel therapeutic strategies that target the underlying mechanisms of PD.

*Hedyotis Diffusae Herba* (HDH), is an annual diffuse herbaceous plant belonging to the Rubiaceae family and the genus Hedyotis. It is also a commonly used traditional Chinese medicinal material. First recorded in Guangxi Chinese Materia Medica Records, it is widely distributed in tropical Asia, stretching from Nepal in the west to Japan in the east. This plant usually grows in moist fields and is not tolerant of drought or waterlogging(R. Chen, He, Tong, Tang, & Liu, 2016).HDH is rich in a variety of active ingredients, such as flavonoids, iridoids, anthraquinones, polysaccharides, and so on. Modern medical research shows that these ingredients possess significant biological activities, including anti-inflammatory, antioxidant, anti-fibroblast, anti-cancer, immunomodulatory, and neuroprotective pharmacological activities (R. Chen et al., 2016).Firstly, in terms of anti-inflammatory effects, the total flavonoids, stigmasterol, β-sitosterol, quercetin, kaempferol, and other extracts play key roles. The total flavonoids can reduce the expression of pro-inflammatory mediators such as nitric oxide (NO), tumor necrosis factor-α (TNF-α), interleukin-6 (IL-6), and interleukin-1β (IL-1β) induced by lipopolysaccharide (LPS) without causing cytotoxicity(Y. Chen, Lin, Li, & Li, 2016). Stigmasterol can inhibit pro-inflammatory mediators such as TNF-α, IL-6, IL-1β, inducible nitric oxide synthase (iNOS), and cyclooxygenase-2 (COX-2), and at the same time, it can increase the production of the anti-inflammatory cytokine IL-10. β-sitosterol can inhibit the proliferation and migration of endothelial cells. Quercetin can activate sirtuin 1 (SIRT1), and then inhibit the release of pro-inflammatory cytokines such as IL-6, TNF-α, IL-1β, interleukin-8 (IL-8), interleukin-13 (IL-13), and interleukin-17 (IL-17). Some studies have indicated that inflammation plays a central role in PD (Liu, Chen, & Chang, 2022; Pajares, A, Manda, Boscá, & Cuadrado, 2020; Simon, Tanner, & Brundin, 2020). These anti-inflammatory properties may theoretically help delay the progression of PD. Secondly, in terms of antioxidant stress, HDH can activate the NRF2/ARE signaling pathway in human prostate tissue and cell models, relieve inflammation and oxidative stress, and prevent and treat chronic prostatitis(Mo, Xia, & Wu, 2024). For PD, oxidative stress is one of the important pathogenic factors. Excessive free radicals attacking structures such as mitochondria can lead to neuronal dysfunction(Simon et al., 2020). The NRF2/ARE signaling pathway is activated by the extracts of HDH can enhance the cell’s antioxidant defense system, scavenge excessive free radicals, and reduce the damage to neurons caused by oxidative stress, thus opening up new ideas for the prevention and treatment of PD(Zgorzynska, Dziedzic, & Walczewska, 2021).Then, its ethanol extracts and other components can also regulate the immune function of the human body, enhance the body’s resistance to pathogens(Kuo et al., 2015), contribute to the recovery of intestinal infections, and have a certain curative effect on rheumatoid arthritis(Deng et al., 2023; J. Jiang et al., 2024). From the perspective of PD, immune imbalance can induce neuroinflammation and exacerbate neuronal damage(Liu et al., 2022). HDH can be used to regulate immune function, which may play a certain role in the prevention and treatment of PD. Some studies have shown that the water extract of HDH can significantly protect the renal tissue and strongly inhibit TNF-α, IL-1β, IL-6, and monocyte chemoattractant protein-1 (MCP-1)(Xu et al., 2022). In addition, five flavonol glycosides and four o-acylated iridoid glycosides were extracted from the MeOH extract of the whole plant of HDH showing significant neuroprotective activity(Kim et al., 2001). It also contains active ingredients mainly composed of iridoids for anti-Alzheimer’s disease(J. Chen et al., 2024), providing new ideas for the remedy of neurodegenerative diseases such as PD.

However, the research about HDH by scholars is also limited, although HDH has potential neuroprotective activity, there is no official quality standard to control the quality. The content of bioactive compounds in HDH varies greatly in samples from different sources and collected at different moments. In addition, there are relatively few related pharmacokinetic studies about HDH, so it’s hard to evaluate its functions to human body. Although there is a large number of published scientific literature on the chemical composition, pharmacological activities and quantitative analysis of HDH, there is no systematic and up-to-date review about it (R. Chen et al., 2016; Kim et al., 2001). Consequently, after recognizing the gaps of existing studies, we propose to employ diversified computational techniques such as network pharmacology, molecular docking and molecular dynamics simulation to achieve deeper comprehension of the pharmacological processes the medications cure the ailments. Network pharmacology is the novel research method combining systems biology, network analysis and pharmacology to explore the multi-target impact of complex natural compounds (Zhao et al., 2023). Molecular docking is the computational simulation technique designed to predict the binding interactions and affinities between little compound drugs and protein receptors (Yu et al., 2023). Molecular dynamics simulation further verifies, optimize the binding modes between the little molecule drugs and the protein receptors, thereby tends to reveal the mechanisms of action more accurately at molecular level (Bai et al., 2023).

Some studies have found that during the development of PD, protein kinase B (PKB/AKT1) may participate in protection against PD through different signaling pathways(Xiromerisiou et al., 2008). IL-6, TNF-α, etc. are potential biomarkers of neuroinflammation in PD(Liu et al., 2022). Tracking these markers in cerebrospinal fluid or blood is helpful for the early diagnosis and monitoring of PD(Liu et al., 2022). At the same time, the body’s response to exogenous stimuli may affect the homeostasis of the nervous system(Y. Jiang et al., 2024; Pajares et al., 2020). As a traditional Chinese medicinal material, HDH has multiple ingredients and abundant targets. However, whether it has therapeutic effect in PD and its mechanism have not been proved and proposed by conclusive research results. Based on this, this study will use network pharmacology, molecular docking,molecular dynamics simulation and cell experiments to analyze and explore how its active ingredients act on key enrichment pathways, clarify its mechanisms of action in neuroprotection, inflammation regulation, etc. and strive to discover new targets and ideas for the treatment of PD (**Figure 1**).

**Figure 1.**
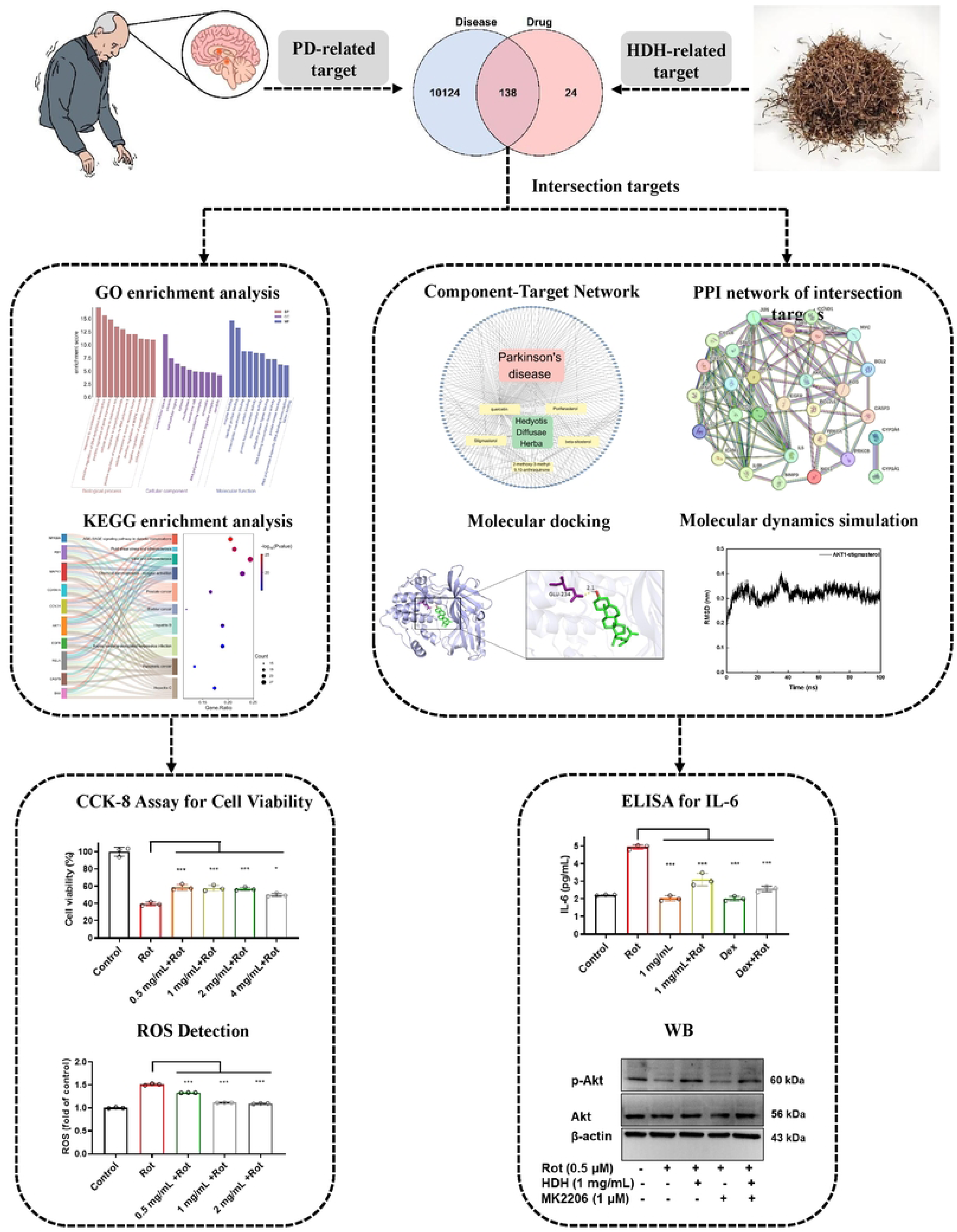
The mechanisms of HDH in anti-PD.

## 2 Materials and Methods

### 2.1 Reagents and Cell Line

HDH extract was procured from Yuanye Bio (Shanghai, China).Human interleukin - 6 (IL - 6) ELISA kit was obtained from Yojianbio (Beijing, China).MK - 2206 2HCl (purity ≥ 98%) was purchased from Macklin Biochemical Co., Ltd. (Shanghai, China).Alisol B was acquired from Afa Biotech (Chengdu, China).Rotenone was obtained from Aladdin Industrial Corporation (Shanghai, China).Reactive oxygen species (ROS) detection kit and Cell Counting Kit - 8 (CCK - 8) were purchased from Beyotime Biotechnology (Shanghai, China).Human neuroblastoma SHSY5Y cells were used in this study.

### 2.2 Cell Culture and Treatment

SH-SY5Y cells were cultured in DMEM/F12 supplemented with fetal bovine serum and antibiotics). Cells were pretreated with HDH extract (0.5, 1, 2, 4 mg/mL) for a certain period, followed by exposure to rotenone (0.5 μM) to establish the injury model. Dexamethasone was used as a positive control. For mechanistic experiments, alisol B (10 μM) and MK2206 (1 μM) were applied as intervention agents.

### 2.3 Screening for Active Components and HDH Targets

The bioactive constituents of HDH were identified by exploring the Traditional Chinese Medicine Systems Pharmacology (TCMSP) (https://old.tcmsp-e.com/tcmsp.php, accessed on 11 January 2025) database. The filtering criteria were oral bioavailability (OB) ≥ 30% and drug-likeness (DL) ≥ 0.18. The structural files of the collected effective components were submitted to the web platform (https://www.novopro.cn/tools/mol2smiles.html, accessed on 17 January 2025) to extract the associated SMILES codes, after which the structures were examined inside the SwissTargetPrediction (http://www.swisstargetpredict.ch/, accessed on 17 January 2025) database. Following an inquiry into the TCMSP database for pharmacological targets, Strawberry Perl software was employed to extract the targets of effective components, converting gene names to gene symbols.

### 2.4 Identification of PD-Related Targets

To forecast PD-related targets, the phrase “Parkinson’s disease” was applied in the GeneCards database (https://www.genecards.org/, accessed on 13 January 2025). R 4.3.2 was subsequently employed to ascertain the overlap between the targets pertinent to HDH and those linked to PD. The VennDiagram package was utilized to create a Venn diagram of the intersecting targets. These overlapping targets served as the effective component targets through which HDH exerts its influence on PD.

### 2.5 Construction of the Component-Target Network

A network was constructed using Cytoscape 3.10.2 software to visualize the connections among the drug, active ingredients, key targets, and disease, specifically for the treatment of PD with HDH.

### 2.6 Protein-Protein Interaction (PPI) Network Analysis

”Homo sapiens” was chosen as the organism after the possible targets of HDH for the treatment of PD were entered into the STRING database (http://www.string-db.org) accessed on 16 January 2025). Concealed free targets were adopted to collect data on protein interactions, with a minimum required interaction score established at 0.9. A total of 30 core targets were found by filtering those having a degree value of ≥ 7. The network of protein interactions for the major targets emerged next.

### 2.7 Gene Ontology (GO) and Kyoto Encyclopedia of Genes and Genomes (KEGG) Pathway Enrichment Analysis

With the settings assigned to Homo sapiens, the PD-related targets were uploaded to the DAVID platform in order to determine their MF, CC, and BP. To create a three-in-one bar chart for GO enrichment analysis, the enrichment findings were shown using the online website (http://www.bioinformatics.com.cn/, accessed on 22 January 2025). The top 10 enriched results were chosen and displayed in a Sankey diagram after KEGG analysis was carried out on the medication- and disease-related target genes using the org.Hs.eg.db and clusterProfiler packages in R.

### 2.8 Molecular Docking

The key target proteins associated with PD were retrieved from the RCSB PDB database (https://www.rcsb.org) with a resolution of 2.5 Å. The chemical structures of the active compounds in HDH were obtained from the TCMSP database and prepared for molecular docking experiments. Molecular docking was performed using AutoDock Tools (version 1.5.7), and binding affinities (docking scores) were calculated. Binding energies lower than -5 kcal/mol were considered indicative of good affinity and stability. The docking results were visualized, and the binding conformations were analyzed and validated using PyMOL software.

### 2.9 Molecular Dynamics Simulation

The Gromacs 2020.3 software was utilized to conduct molecular dynamics simulations to assess the binding stability between proteins and traditional Chinese medicine (TCM) molecules. The molecular docking results were imported into the software, with the Amber99sb-lidn force field selected for the protein molecules and the GAFF force field for the small molecules. The protein-TCM molecule complex was then placed in a TIP3P water model, and parameters were adjusted to position the protein-TCM molecule complex at the center of a cubic simulation box with a periodic boundary of 1.2 nm. The system parameters were set to 0.9% NaCl, a pH of 7.0, a temperature of 300 K, and a pressure of one atmosphere to simulate the internal human environment. After the setup, the system’s energy was minimized, and a 100 ns molecular dynamics simulation with a time step of 2 fs was performed on the complex. Structural coordinates were saved every 100 ps. the root mean square deviation (RMSD), root mean square fluctuation (RMSF), and radius of gyration (Rg) were analyzed to evaluate the stability of the complexes.

### 2.10 CCK-8 Assay for Cell Viability

After treatment, CCK-8 solution was added to each well. After incubation, the absorbance at 450 nm was measured using a microplate reader. Cell viability was calculated relative to the control group.

### 2.11 ROS Detection

Intracellular ROS levels were detected using the ROS assay kit following the manufacturer’s instructions. Fluorescence intensity was measured, and the results were expressed as fold change relative to the control.

### 2.12 ELISA for IL-6

Cell culture supernatants were collected, and IL-6 levels were determined using the human IL-6 ELISA kit according to the protocol provided by Yojianbio.

### 2.13 Western Blot Analysis

Total protein was extracted, and the expression levels of p-Akt, Akt, and β-actin were detected by Western blot. The relative density of p-Akt to Akt was analyzed using image analysis software.

### 2.14 Statistical Analysis

Data were presented as mean ± standard deviation (SD). Statistical comparisons were performed using one-way ANOVA followed by post hoc tests. Statistical significance is denoted as *** *p* < 0.001, ** *p* < 0.01, * *p* < 0.05.

## 3 Results

### 3.1 Active Components and Predicted Targets of HDH

From the TCMSP database, seven active components of HDH were found using the filtering criteria of oral bioavailability ≥ 30% and drug-likeness > 0.18 (**Table 1**). The architecture of the seven effective components derived from the SwissTargetPrediction database (**Figure 2**), including 2,3-dimethoxy-6-methyanthraquinone, 2-methoxy-3-methyl-9,10-anthraquinone, quercetin, (4aS,6aR,6aS,6bR,8aR,10R,12aR,14bS)-10-hydroxy-2,2,6a,6b,9,9,12a-heptamethyl-1,3,4,5,6,6a,7,8,8a,10,11,12,13,14b-tetradecahydropicene-4a-carboxylic acid, stigmasterol, β-sitosterol and quercetin. A total of 556 drug targets were obtained through TCMSP, which were subsequently filtered and deduplicated to yield 162 targets corresponding to the effective active compounds.

**Table 1.**
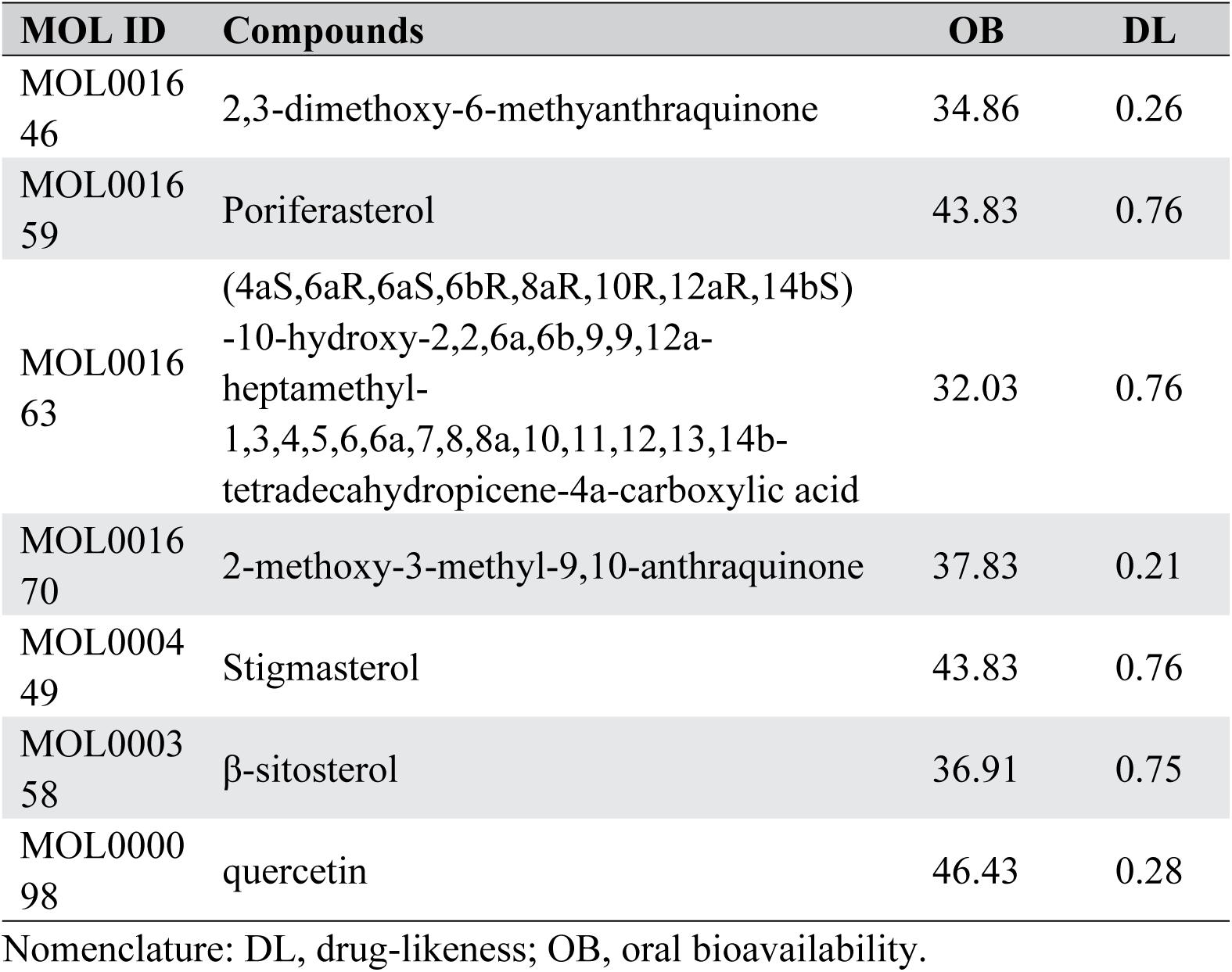
Screening Results of Active Compounds of HDH Identified by TCMSP.

**Figure 2.**
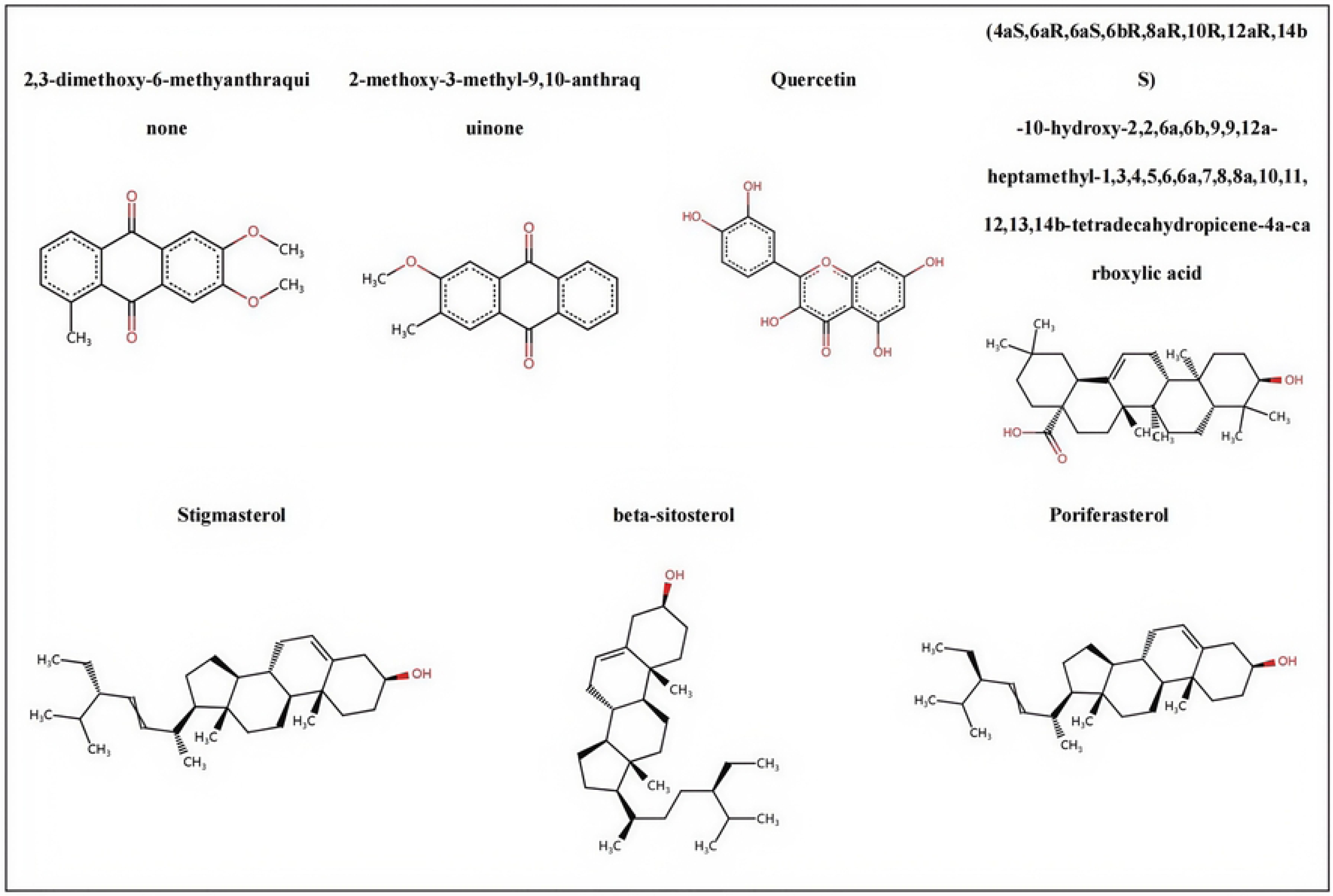
Structural formulas of seven HDH components.

### 3.2 PD-Related targets

A total of 10,262 disease targets were obtained by gathering PD-related targets from the GeneCards database. The targets of the PD-related targets and the active components of HDH were intersected to form the Venn diagram shown in **Figure 3A**, which produced 138 intersecting targets in total.

**Figure 3.**
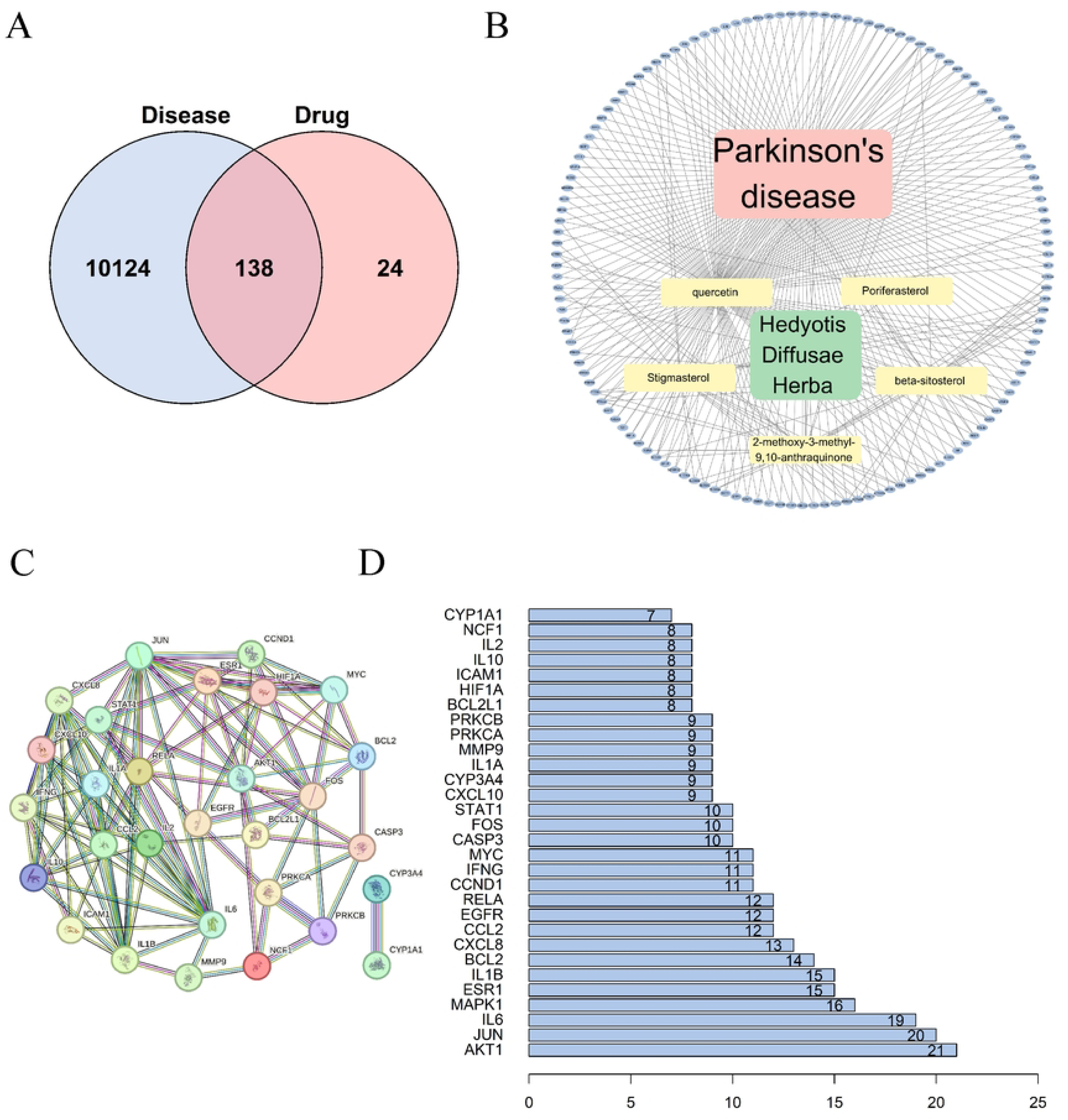
**(A)** Shared Molecular Targets of HDH and PD. The diagram illustrates 10,262 PD-related targets (left), targets of active ingredients of HDH (right), and 138 common molecular targets (center). **(B)** Target network of bioactive compounds from HDH in PD. The constructed network comprises 145 nodes, including 1 drug, 1 disease, 5 bioactive compounds, and 138 targets connected by 327 edges. **(C)** Construction of the PPI Network for Targets Regulated by HDH. This network comprises 29 nodes and 105 edges, with an average node degree 7.24. These nodes represent potential targets implicated in PD. **(D)** Degree Distribution of Core Genes in the PPI Network. The degree distribution analysis highlights key proteins’ central roles and functional importance within the PPI network.

### 3.3 Traditional Chinese medicine regulatory network

Data on HDH’s active ingredients and targets was imported into Cytoscape 3.10.2 to create the regulatory network for traditional Chinese medicine. This network (**Figure 3B**) has 327 edges and 145 nodes, including 138 core target nodes, five active component nodes, one medicine node, and one illness node. According to the results, several HDH constituents, including quercetin, β-sitosterol, 2-methoxy-3-methyl-9,10-anthraquinone, stigmasterol, and poriferasterol, can be used to identify appropriate targets for PD treatment.

### 3.4 PPI network and core targets

The PPI network analysis identified 29 core targets, with AKT1, JUN, and IL-6 being the most central nodes (**Figure 3C**). These targets are known to play critical roles in neuroinflammation and oxidative stress, which are key pathological mechanisms in PD. In addition, the top thirty core targets are presented in bar chart (**Figure 3D**).

### 3.5 GO and KEGG Pathway Enrichment Analysis

GO and KEGG enrichment analyses were conducted on the pertinent targets in order to investigate the biological roles and signaling pathways of important genes. The GO BP included 577 entries, mostly pertaining to response to xenobiotic stimulus, positive regulation of DNA-templated transcription, and positive regulation of gene expression. Some like extracellular space, caveola, and extracellular region were involved in the CC, which had a cellular section of 71. In addition, 144 were for the MF, including enzyme binding, identical protein binding, transcription coactivator binding, etc. The top ten biological functions were chosen and shown in a combined bar chart (**Figure 4A**), where smaller P values denoted more relevance. After 168 KEGG pathways were enriched using a filtering parameter of p < 0.05, the top 10 pathways with the highest enrichment were chosen to produce a Sankey diagram (**Figure 4B**), which included pathways like lipid and atherosclerosis, chemical carcinogenesis-receptor activation, and fluid shear stress and atherosclerosis.

**Figure 4.**
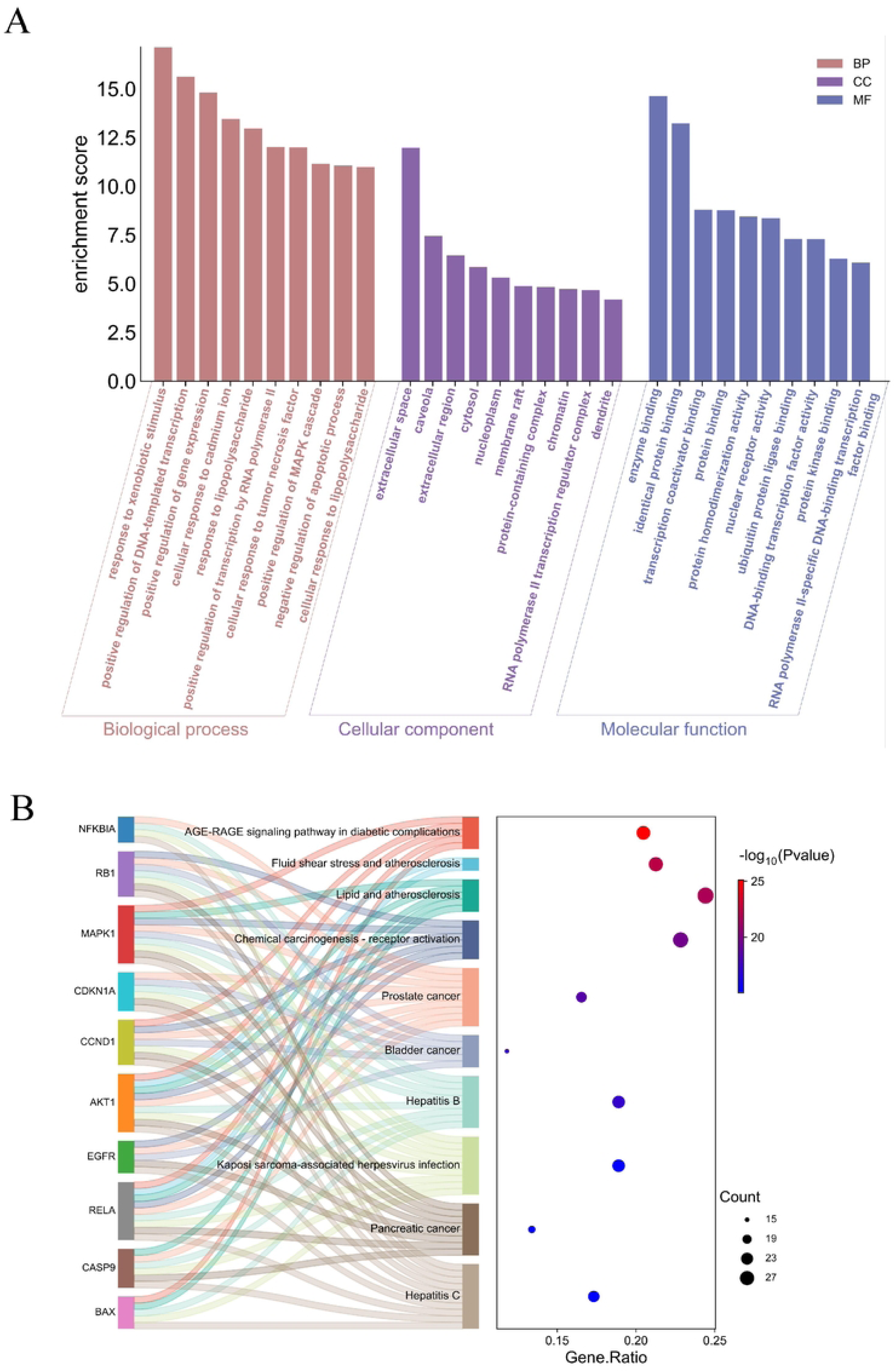
**(A)** GO Enrichment of HDH Targets in PD. The top 10 enriched biological processes (BP), cellular components (CC), and molecular functions (MF) are illustrated in a bar chart. The x-axis represents the terms, while the y-axis indicates fold enrichment. BP, CC, and MF are depicted in rosy brown, purple, and slate blue, respectively. **(B)** An enrichment analysis based on KEGG pathways was conducted for the identified targets. A Sankey bubble diagram visualizing the top 10 significantly enriched pathways (P < 0.05) is provided.

### 3.6 Molecular Docking Verification

The binding capacities of five core active components from HDH (quercetin, stigmasterol, β-sitosterol, poriferasterol, and 2-methoxy-3-methyl-9,10-anthraquinone) with PD-related targets (AKT1, IL-6, and JUN) were evaluated through molecular docking simulations, using levodopa as a positive control. As shown in **Table 2**, nearly all tested complexes exhibited favorable binding energies below -5 kcal/mol. Notably, the binding energies of HDH components for all three targets were significantly lower than those of levodopa, indicating superior binding affinities. Figure 5 presents the optimal docking conformations for three representative interactions. Specifically, stigmasterol demonstrated the strongest binding to AKT1 (ΔG = -8.52 kcal/mol) through insertion of its steroid nucleus into the active pocket. A hydrogen bond (2.1 Å) formed between the hydroxyl group of stigmasterol and GLU-234 residue (**Figure 5A**), suggesting potential competitive inhibition of AKT1 kinase activity that might activate downstream anti-apoptotic pathways to protect dopaminergic neurons. For JUN protein, poriferasterol showed optimal binding (ΔG = -7.82 kcal/mol) via hydrogen bonding (2.3 Å) between its hydroxyl group and GLU-552 residue (**Figure 5B**). β-sitosterol exhibited the highest affinity for IL-6 (ΔG = -8.62 kcal/mol), forming a strong hydrogen bond (2.0 Å) with GLY-59 residue in the binding pocket (**Figure 5C**). The findings suggest that natural compounds from HDH may achieve enhanced therapeutic efficacy through superior target occupation compared to conventional medications of PD, potentially providing a theoretical foundation for developing complementary treatment strategies.

**Table 2.** Molecular docking results of HDH components with core targets.

| Compound<br>Name | Affinity (kcal/mol) |  |  |  |
| --- | --- | --- | --- | --- |
|  | AKT1 | IL-6 | JUN | Mean |
| Quercetin | -5.99 | -6.36 | -5.84 | -6.06 |
| Poriferasterol | -8.01 | -6.8 | -7.82 | -7.54 |
| Stigmasterol | -8.52 | -7.89 | -7.42 | -7.94 |
| β-sitosterol | -7.78 | -8.62 | -7.44 | -7.95 |
| 2-methoxy-3-methyl-9,10-anthraquinone | -7.46 | -6.59 | -7.1 | -7.05 |
| Mean | -7.55 | -7.25 | -7.12 |  |
| Levodopa | -5.73 | -5.95 | -5.33 | -5.71 |

**Figure 5.**
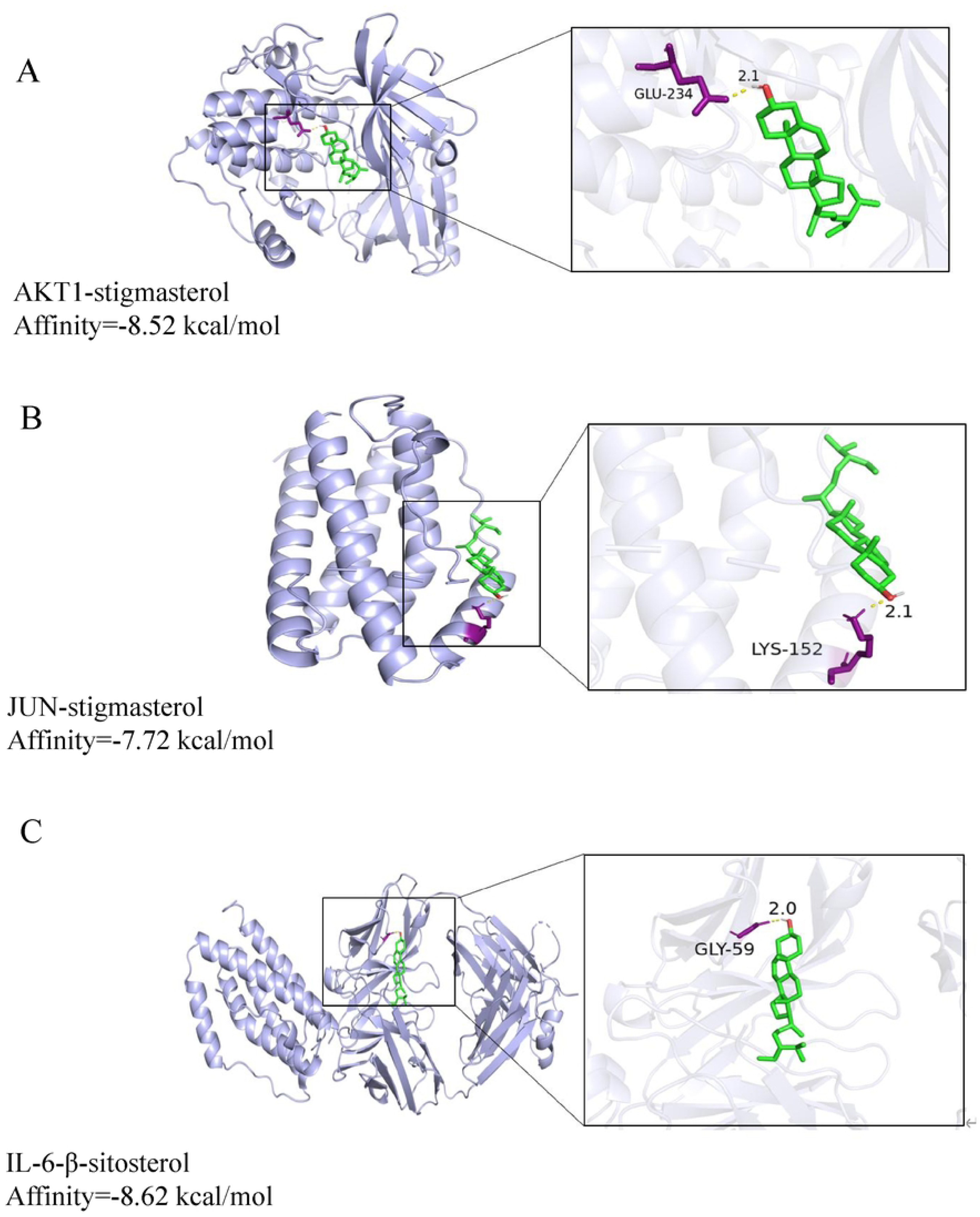
Molecular Docking of HDH Active Compounds with Target Proteins in PD. **(A)** AKT1-stigmasterol. **(B)** JUN-stigmasterol. **(C)** IL-6-β-sitosterol.

### 3.7 Molecular Dynamics Simulations (MDS) and Binding Free Energy Calculations

To investigate the interactions between receptor proteins and small molecules during dynamic processes and evaluate binding site stability, we performed 100 ns molecular dynamics simulations on the complexes. System equilibration was validated through root mean square deviation (RMSD) analysis, a critical metric for assessing system stability. As shown in **Figure 6A**, the AKT1-stigmasterol, JUN-poriferasterol, and IL-6-β-sitosterol systems reached equilibrium states after 5, 40, and 50 ns of simulation, respectively. The final RMSD ranges stabilized at 0.2601–0.4107 nm for AKT1-stigmasterol, 0.8235–1.0418 nm for JUN-poriferasterol, and 0.5086–0.5580 nm for IL-6-β-sitosterol, with AKT1-stigmasterol exhibiting the smallest fluctuations. Radius of gyration (Rg) analysis demonstrated progressive structural compaction during simulations (**Figure 6B**). Final Rg values stabilized at 2.17–2.25 nm (AKT1-stigmasterol), 1.85–2.09 nm (JUN-poriferasterol), and 2.97–3.08 nm (IL-6-β-sitosterol), suggesting enhanced structural density and flexibility. The converging Rg profiles imply comparable structural compactness between systems.Solvent-accessible surface area (SASA) measurements (**Figure 6C**) stabilized at 190, 229, and 257 nm² for the respective complexes, confirming the critical role of hydrophobic core formation in maintaining structural stability.

**Figure 6.**
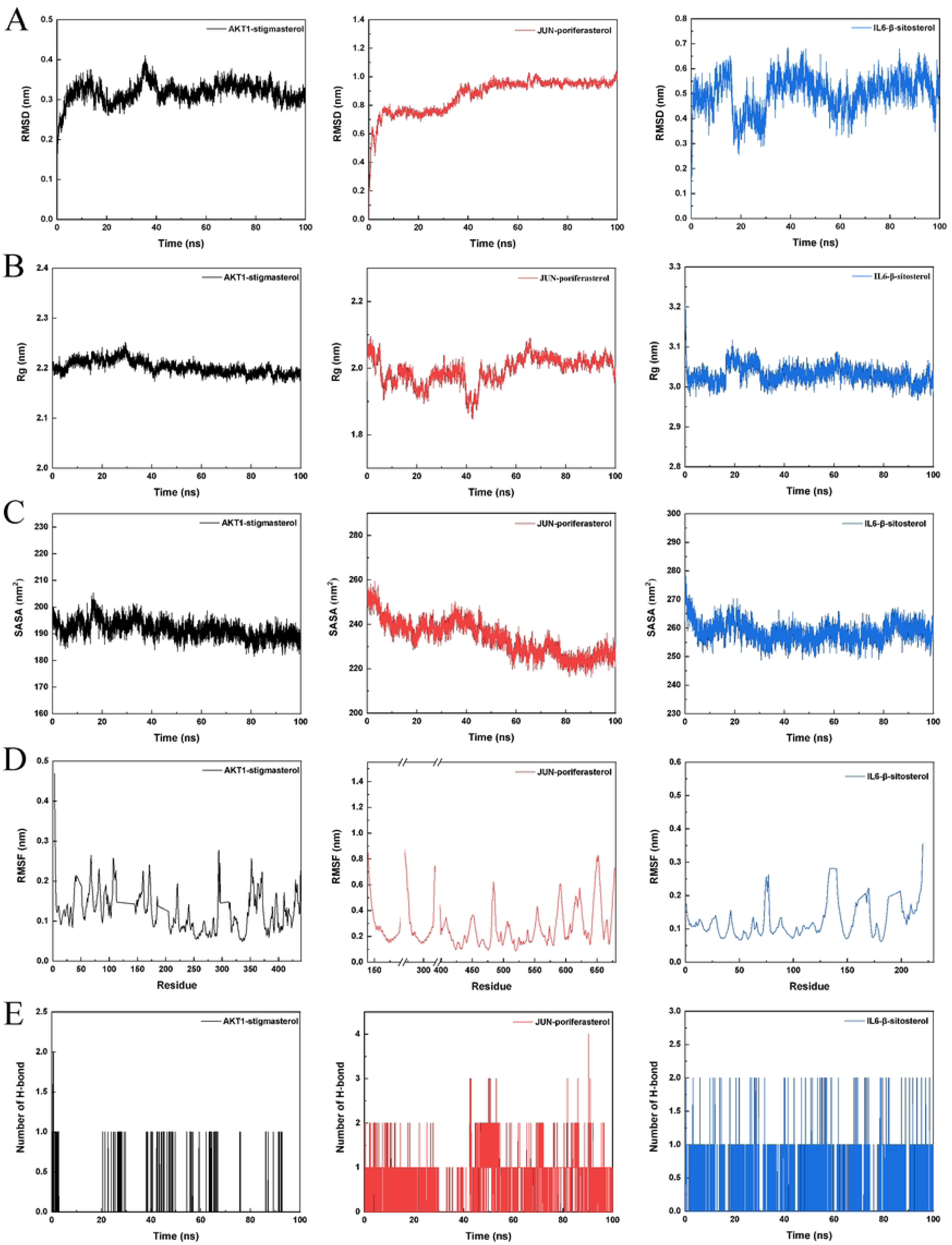
**(A)** Root Mean Square Deviation (RMSD) values of three complexes. **(B)** Radius of Gyration (Rg) values of three complexes. **(C)** Solvent-accessible surface area (SASA) values of three complexes. **(D)** Root Mean Square Fluctuation (RMSF) values of the three complexes. **(E)** The number of hydrogen bonds in three complexes.

Root mean square fluctuation (RMSF) analysis revealed residue-specific flexibility across the complexes (**Figure 6D**). While both systems displayed similar RMSF trends, minor variations (<0.6 nm) in specific residues were attributed to differential binding interactions. Notably, all RMSF values remained below 0.6 nm, indicating exceptional protein rigidity and dynamic stability—a finding consistent with high-stability systems reported in prior studies.Hydrogen bond analysis revealed distinct interaction patterns **(Figure 6E)**. During simulations, average hydrogen bond counts were 0.02 (AKT1-stigmasterol), 0.61 (JUN-poriferasterol), and 0.34 (IL-6-β-sitosterol), with JUN-poriferasterol showing the strongest hydrogen bonding network.

Binding energy calculations (**Table 3**) identified AKT1-stigmasterol as the most stable complex, with superior binding energy compared to JUN-poriferasterol and IL-6-β-sitosterol. Reduced binding affinities in the latter systems correlated with increased conformational flexibility and torsional angles near key binding residues. Energy decomposition analysis (**Figure 7**) highlighted residue-specific contributions to binding free energy. AKT1-stigmasterol exhibited more negative decomposition values at active site residues than other complexes, indicating stronger interactions and superior complementarity. This mechanistic insight aligns with its enhanced binding affinity observed in earlier analyses.

**Table 3.** Binding Free Energies and Energy Contributions for AKT1-stigmasterol, JUN-poriferasterol, and IL-6-β-stiosterol Complexes.

| No. | Electrostatic<br>energy<br>(KJ<br>mol<sup>-1</sup>) | Van der<br>Waals<br>Energy<br>(KJ<br>mol<sup>-1</sup>) | Polar<br>solvation<br>energy<br>(KJ<br>mol<sup>-1</sup>) | Nonpolar<br>solvation<br>Energy<br>(KJ<br>mol<sup>-1</sup>) | Total<br>Binding<br>Energy<br>(KJ<br>mol<sup>-1</sup>) |
| --- | --- | --- | --- | --- | --- |
| AKT1-stigmasterol | -2.575 | -190.761 | 46.922 | -23.577 | -169.992 |
| JUN-poriferasterol | -15.766 | -148.249 | 78.490 | -22.154 | -107.679 |
| IL-6-β-<br>sitosterol | -22.037 | -178.540 | 138.918 | -23.643 | -85.302 |

**Figure 7.**
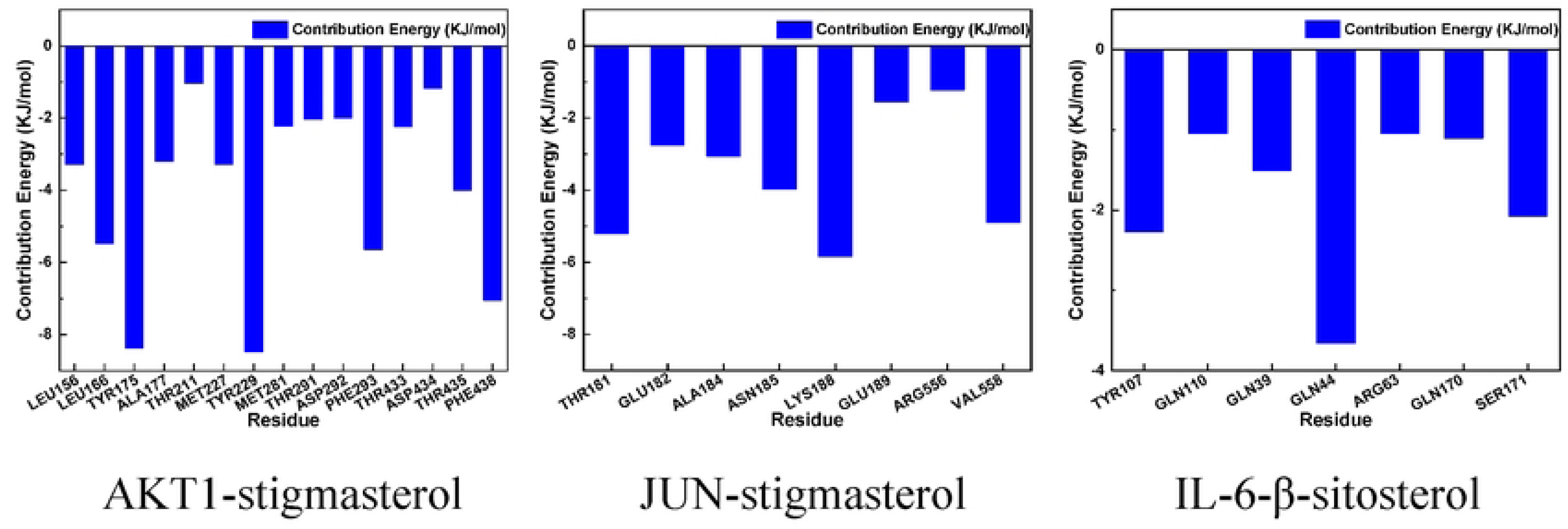
Energy contributions of key amino acid residues in three complexes. From left to right are: AKT1-stigmasterol, JUN-poriferasterol, IL-6-β-sitosterol.

### 3.8 HDH Extract Alleviates Rotenone - Induced Impairments in Cell Viability and ROS Homeostasis

Rotenone (0.5 μM) exerted a profound cytotoxic effect, significantly decreasing SH - SY5Y cell viability when compared to the control group (**Figure 8A**). Conversely, HDH extract, within the concentration range of 0.5–4 mg/mL, enhanced cell viability in rotenone - treated cells. Marked protective effects were observed at concentrations of 1–4 mg/mL, as evidenced by significant differences compared to the Rot group. Parallel to its impact on cell viability, rotenone exposure triggered a substantial surge in intracellular ROS levels (**Figure 8B**). Notably, HDH extract at concentrations of 0.5 and 1 mg/mL effectively mitigated these elevated ROS levels in rotenone - injured cells, thereby implying the extract’s inherent antioxidant properties.

**Figure 8.**
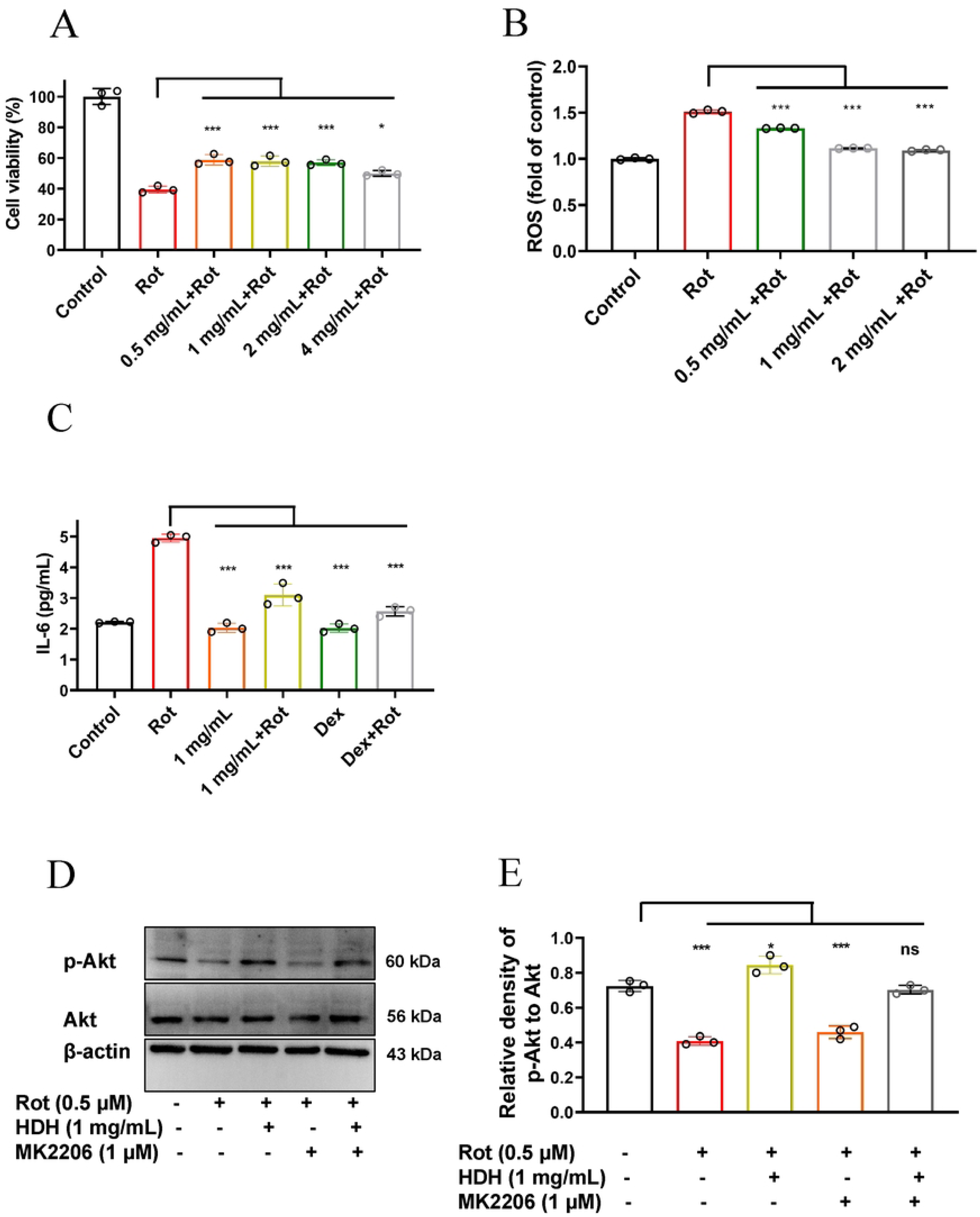
Protective effects of HDH against Rot-induced injury in SH-SY5Y cells.**(A)** Cell viability analysis via CCK-8 assay.SH-SY5Y cells were treated with Rot (0.5 μM) alone or pre-treated with HDH (0.5–4 mg/mL) prior to Rot exposure.**(B)** Intracellular reactive oxygen species (ROS) measurement. **(C)** IL-6 secretion analyzed by ELISA. **(D)** Western blot analysis of p-Akt and Akt.**(E)** The bar chart indicates the normalization of the level of p-Akt to total Akt.Data are mean ± SD (n ≥ 3). *** *p* < 0.001, ** *p* < 0.01, * *p* < 0.05.

### 3.9 HDH Extract Modulates Inflammatory Response and Akt Signaling Pathway

Rotenone stimulation resulted in a substantial increase in the secretion of interleukin-6 (IL-6) in SH - SY5Y cells compared to the control (**Figure 8C**). Pretreatment with HDH (1 mg/mL) significantly reduced IL-6 levels, an effect comparable to that of dexamethasone (Dex, positive control), suggesting the anti - inflammatory property of HDH.Western blot analysis (**Figure 8D** and **8E**) showed that rotenone treatment decreased the phosphorylation level of Akt (p - Akt/Akt ratio) compared to the control. HDH (1 mg/mL) intervention enhanced the phosphorylation of Akt in rotenone - exposed cells. However, when MK2206 (an Akt inhibitor) was co - administered, the promoting effect of HDH on Akt phosphorylation was reversed, indicating that HDH might exert its protective effects through the Akt signaling pathway.

## 4. Discussion

PD is a debilitating neurodegenerative disorder that poses a significant global health burden (Poewe et al., 2017). The disease is characterized by the progressive loss of dopaminergic neurons in the substantia nigra and the pathological accumulation of α-synuclein, leading to motor symptoms such as bradykinesia, rigidity, and tremors, as well as non-motor manifestations including cognitive decline and autonomic dysfunction. Current therapeutic strategies, primarily centered on dopamine replacement, provide symptomatic relief but fail to halt disease progression and are often accompanied by severe side effects with prolonged use(Olanow, Stern, & Sethi, 2009). This underscores the critical need for novel, multi-targeted therapeutic approaches. HDH, a traditional Chinese medicinal herb with demonstrated anti-inflammatory, antioxidant, antitumor and immunomodulatory effects, emerges as a promising candidate for PD treatment(Huang et al., 2022; Hung et al., 2022; Q. Li et al., 2018).

Chronic inflammation is considered a significant cause of PD, and reducing inflammation may aid in the prevention and treatment of PD(Bottigliengo et al., 2022; Zhang, Xiao, Mao, & Xia, 2023). Consequently, five key HDH components were further analyzed for their therapeutic effects on PD. Network pharmacology analysis revealed that the bioactive compounds in HDH, such as quercetin, poriferasterol, stigmasterol,β-sitosterol and 2-methoxy-3-methyl-9,10-anthraquinone, exert their therapeutic effects by modulating multiple signaling pathways, including PI3K/AKT, MAPK, and NF-κB in several diseases(Lv, Pan, Song, Chang, & Zhang, 2021; Mo et al., 2024; Ye et al., 2024). Stigmasterol and β-sitosterol are two major phytosterols present in HDH. Stigmasterol suppresses the inflammatory response and is involved in the treatment of Alzheimer’s disease through the NF-κB and NLRP3 signaling pathways (Jie et al., 2022). β - sitosterol, on the other hand, inhibits neuroinflammation by impeding the TLR4/NF - κB signaling pathway (Zheng et al., 2023).Furthermore, certain components in HDH can traverse the blood-brain barrier and act directly on the central nervous system, thereby enhancing its therapeutic potential (Jie et al., 2022). These findings provide a theoretical foundation for the application of HDH in the treatment of PD. The synergistic effects of these components suggest that HDH holds the potential for multi-target therapy in PD treatment, capable of decelerating neuronal degeneration and death via multiple pathways.

PPI network analysis has revealed three potential key targets—AKT1, JUN, and IL-6—that are closely associated with the treatment of PD using HDH. These genes may play a role in the therapeutic mechanisms by which HDH alleviates PD. It has been demonstrated that genetic variations in AKT1 can affect brain function in individuals with PD (X. Y. Li et al., 2016). The JUN gene encodes the protein c-Jun, a transcription factor (Eferl & Wagner, 2003; Shaulian & Karin, 2002). In PD, neuroinflammation is a critical pathological mechanism, and c-Jun plays a key role in inflammatory responses, potentially influencing disease progression by regulating the expression of inflammatory mediators [17-19]. Numerous clinical studies have found that levels of IL-6 in the blood and cerebrospinal fluid of PD patients are significantly higher than those in healthy individuals. Moreover, changes in IL-6 levels are correlated with the severity and progression of PD (Milyukhina, Karpenko, & Klimenko, 2015; Qu et al., 2023). Therefore, AKT1, JUN, and IL-6 are closely related to the onset and development of PD, and the components of HDH may exert their anti-PD effects by interacting with these target proteins.

Molecular docking studies provided further insights into the interactions between the active components of HDH and key PD-related molecular targets. For instance, stigmasterol demonstrated a strong binding affinity for AKT1, a critical kinase in the PI3K/AKT pathway, which is known to promote neuronal survival and inhibit apoptosis. Similarly, β-sitosterol exhibited high binding affinity for IL-6, suggesting its potential to inhibit inflammation of PD [17-19]. These findings not only validate the predictions from network pharmacology but also offer a mechanistic basis for the neuroprotective effects of HDH. Molecular dynamics simulations further reinforced the molecular docking results by elucidating the dynamic stability and conformational changes associated with the binding of HDH compounds to their target proteins. For example, the interaction between AKT1-stigmasterol remained stable over a 100-nanosecond simulation, indicating a robust and energetically favorable binding. Additionally, stigmasterol binding to AKT1 induced conformational changes that enhanced the kinase’s activity, further supporting its role in neuroprotection. These simulations provide a detailed understanding of the molecular interactions underlying the therapeutic potential of HDH.

This study highlights the neuroprotective potential of HDH against rotenone-induced cytotoxicity in SH-SY5Y neuroblastoma cells. The protective mechanisms were associated with enhanced cell viability, reduced oxidative stress and inflammation, and the activation of the Akt signaling pathway. Rotenone, a potent mitochondrial complex I inhibitor, is widely used to model neurodegeneration due to its ability to increase intracellular reactive oxygen species (ROS) and trigger inflammatory cascades such as IL-6 secretion—processes relevant to PD and related disorders (Rahul & Siddique, 2021; Vasudevan Sajini, Thaggikuppe Krishnamurthy, Chakkittukandiyil, & Mudavath, 2024). In line with previous studies, our results confirm that rotenone reduces cell viability and elevates ROS and IL-6 levels in SH-SY5Y cells.HDH treatment significantly attenuated these deleterious effects. Its ability to suppress ROS generation is likely attributable to phytochemicals like scopoletin, which have been shown to modulate oxidative stress and inflammatory mediators such as IL-6 (Gao, Li, Zhang, & Bai, 2024). Additionally, other plant-derived compounds like apigenin and chrysin demonstrate similar protective effects through antioxidant and pathways(Del Fabbro, Bortolotto, Ferreira, Sari, & Furian, 2025; Kashyap, Shikha, Thakur, & Aneja, 2022; Kurt et al., 2023; Singh et al., 2023). Mechanistically, this study shows that HDH reverses rotenone-induced suppression of Akt phosphorylation, implying a critical role for Akt signaling in mediating neuroprotection. The use of MK2206, a selective Akt inhibitor, validated this pathway by abrogating HDH’s protective effects—echoing findings from other studies that underscore the neuroprotective relevance of Akt activation in oxidative stress contexts(Rahul & Siddique, 2021; Singh et al., 2023).Despite these promising results, the study has limitations. The exact bioactive constituents in HDH remain unidentified. HDH is rich in flavonoids, saponins, and polysaccharides—each with potential neuroprotective activities that warrant further isolation and characterization. Furthermore, while *in vitro* studies are informative, *in vivo* validation in animal models is essential for clinical translation.

## 5. Conclusion

This study systematically elucidates the neuroprotective mechanisms of HDH against rotenone-induced cytotoxicity in SH-SY5Y cells by integrating network pharmacology, molecular simulations, and experimental validation. Key bioactive compounds such as stigmasterol, quercetin, and β-sitosterol exhibited strong affinities with core PD-related targets including AKT1, IL-6, and JUN, as confirmed by molecular docking and dynamics simulations. Functionally, HDH extract restored cell viability, reduced intracellular ROS accumulation, and suppressed IL-6 secretion in rotenone-injured neurons. Moreover, HDH significantly upregulated Akt phosphorylation, and this effect was reversed by the Akt-specific inhibitor MK2206, indicating that the Akt pathway is critically involved in mediating its protective effects. Collectively, these findings provide both mechanistic and experimental support for HDH as a promising multi-target candidate for the prevention and adjunctive treatment of PD.

## Data availability

Data will be made available on request.

## Acknowledgements

This research was funded by grants from Young Innovation Project of Sichuan Medical Association (No. Q2024077), Scientific Research Fund of the Affiliated Hospital of Southwest Medical University (24234), and Project of the Luzhou Municipal Science and Technology Bureau (Project No: 2022-JYJ-152). Artificial intelligence tools were used to assist with language polishing during manuscript preparation.

## References

Bai, G., Pan, Y., Zhang, Y., Li, Y., Wang, J., Wang, Y., … Cao, J. (2023). Research advances of molecular docking and molecular dynamic simulation in recognizing interaction between muscle proteins and exogenous additives. Food Chem, 429, 136836. doi:10.1016/j.foodchem.2023.136836

Bottigliengo, D., Foco, L., Seibler, P., Klein, C., König, I. R., & Del Greco, M. F. (2022). A Mendelian randomization study investigating the causal role of inflammation on Parkinson’s disease. Brain, 145(10), 3444–3453. doi:10.1093/brain/awac193

Chen, J., Rao, J., Lu, H., Lu, M., Wang, C., & Cao, Y. (2024). Network pharmacology and experimental verification to explore the effect of Hedyotis diffusa on Alzheimer’s disease. Chem Biol Drug Des, 103(6), e14558. doi:10.1111/cbdd.14558

Chen, R., He, J., Tong, X., Tang, L., & Liu, M. (2016). The Hedyotis diffusa Willd. (Rubiaceae): A Review on Phytochemistry, Pharmacology, Quality Control and Pharmacokinetics. Molecules, 21(6). doi:10.3390/molecules21060710

Chen, Y., Lin, Y., Li, Y., & Li, C. (2016). Total flavonoids of Hedyotis diffusa Willd inhibit inflammatory responses in LPS-activated macrophages via suppression of the NF-κB and MAPK signaling pathways. Exp Ther Med, 11(3), 1116–1122. doi:10.3892/etm.2015.2963

Del Fabbro, L., Bortolotto, V. C., Ferreira, L. M., Sari, M. H. M., & Furian, A. F. (2025). Chrysin’s anti-inflammatory action in the central nervous system: A scoping review and an evidence-gap mapping of its mechanisms. Eur J Pharmacol, 997, 177602. doi:10.1016/j.ejphar.2025.177602

Deng, H., Jiang, J., Zhang, S., Wu, L., Zhang, Q., & Sun, W. (2023). Network pharmacology and experimental validation to identify the potential mechanism of Hedyotis diffusa Willd against rheumatoid arthritis. Sci Rep, 13(1), 1425. doi:10.1038/s41598-022-25579-3

Eferl, R., & Wagner, E. F. (2003). AP-1: a double-edged sword in tumorigenesis. Nat Rev Cancer, 3(11), 859–868. doi:10.1038/nrc1209

Gao, X. Y., Li, X. Y., Zhang, C. Y., & Bai, C. Y. (2024). Scopoletin: a review of its pharmacology, pharmacokinetics, and toxicity. Front Pharmacol, 15, 1268464. doi:10.3389/fphar.2024.1268464

Huang, F., Pang, J., Xu, L., Niu, W., Zhang, Y., Li, S., & Li, X. (2022). Hedyotis diffusa injection induces ferroptosis via the Bax/Bcl2/VDAC2/3 axis in lung adenocarcinoma. Phytomedicine, 104, 154319. doi:10.1016/j.phymed.2022.154319

Hung, H. Y., Cheng, K. C., Kuo, P. C., Chen, I. T., Li, Y. C., Hwang, T. L., … Wu, T. S. (2022). Chemical Constituents of Hedyotis diffusa and Their Anti-Inflammatory Bioactivities. Antioxidants (Basel), 11(2). doi:10.3390/antiox11020335

Jankovic, J. (2008). Parkinson’s disease: clinical features and diagnosis. J Neurol Neurosurg Psychiatry, 79(4), 368–376. doi:10.1136/jnnp.2007.131045

Jiang, J., Huang, M., Zhang, S. S., Wu, Y. G., Li, X. L., Deng, H., … Sun, W. K. (2024). Identification of Hedyotis diffusa Willd-specific mRNA-miRNA-lncRNA network in rheumatoid arthritis based on network pharmacology, bioinformatics analysis, and experimental verification. Sci Rep, 14(1), 6291. doi:10.1038/s41598-024-56880-y

Jiang, Y., Wu, W., Xie, L., Zhou, Y., Yang, K., Wu, D., … Ge, J. (2024). Molecular targets and mechanisms of Sijunzi decoction in the treatment of Parkinson’s disease: evidence from network pharmacology, molecular docking, molecular dynamics simulation, and experimental validation. Front Pharmacol, 15, 1487474. doi:10.3389/fphar.2024.1487474

Jie, F., Yang, X., Yang, B., Liu, Y., Wu, L., & Lu, B. (2022). Stigmasterol attenuates inflammatory response of microglia via NF-κ B and NLRP3 signaling by AMPK activation. Biomed Pharmacother, 153, 113317. doi:10.1016/j.biopha.2022.113317

Kashyap, P., Shikha, D., Thakur, M., & Aneja, A. (2022). Functionality of apigenin as a potent antioxidant with emphasis on bioavailability, metabolism, action mechanism and in vitro and in vivo studies: A review. J Food Biochem, 46(4), e13950. doi:10.1111/jfbc.13950

Kim, Y., Park, E. J., Kim, J., Kim, Y., Kim, S. R., & Kim, Y. Y. (2001). Neuroprotective constituents from Hedyotis diffusa. J Nat Prod, 64(1), 75–78. doi:10.1021/np000327d

Kuo, Y. J., Lin, J. P., Hsiao, Y. T., Chou, G. L., Tsai, Y. H., Chiang, S. Y., … Chung, J. G. (2015). Ethanol Extract of Hedyotis diffusa Willd Affects Immune Responses in Normal Balb/c Mice In Vivo. In Vivo, 29(4), 453–460.

Kurt, G. A., Ertekin, T., Atay, E., Bilir, A., Koca, H. B., Aslan, E., & Sar ı ta ş, A. (2023). Investigation of the antioxidant effect of Chrysin in an experimental cataract model created in chick embryos. Mol Vis, 29, 245–255.

Li, Q., Lai, Z., Yan, Z., Peng, J., Jin, Y., Wei, L., & Lin, J. (2018). Hedyotis diffusa Willd inhibits proliferation and induces apoptosis of 5 - FU resistant colorectal cancer cells by regulating the PI3K/AKT signaling pathway. Mol Med Rep, 17(1), 358–365. doi:10.3892/mmr.2017.7903

Li, X. Y., Teng, J. J., Liu, Y., Wu, Y. B., Zheng, Y., & Xie, A. M. (2016). Association of AKT1 gene polymorphisms with sporadic Parkinson’s disease in Chinese Han population. Neurosci Lett, 629, 38–42. doi:10.1016/j.neulet.2016.06.052

Liu, T. W., Chen, C. M., & Chang, K. H. (2022). Biomarker of Neuroinflammation in Parkinson’s Disease. Int J Mol Sci, 23(8). doi:10.3390/ijms23084148

Lv, Y. X., Pan, H. R., Song, X. Y., Chang, Q. Q., & Zhang, D. D. (2021). Hedyotis diffusa plus Scutellaria barbata Suppress the Growth of Non-Small-Cell Lung Cancer via NLRP3/NF-κ B/MAPK Signaling Pathways. Evid Based Complement Alternat Med, 2021, 6666499. doi:10.1155/2021/6666499

Milyukhina, I. V., Karpenko, M. N., & Klimenko, V. M. (2015). [Clinical parameters and the level of certain cytokines in blood and cerebrospinal fluid of patients with Parkinson’s disease]. Klin Med (Mosk), 93(1), 51–55.

Mo, J., Xia, K., & Wu, C. (2024). Hedyotis diffusa Willd inhibits inflammation and oxidative stress to protect against chronic prostatitis via the NRF2/ARE signaling pathway. Environ Toxicol, 39(8), 4221–4230. doi:10.1002/tox.24298

Olanow, C. W., Stern, M. B., & Sethi, K. (2009). The scientific and clinical basis for the treatment of Parkinson disease (2009). Neurology, 72(21 Suppl 4), S1–136. doi:10.1212/WNL.0b013e3181a1d44c

Pajares, M., A, I. R., Manda, G., Boscá, L., & Cuadrado, A. (2020). Inflammation in Parkinson’s Disease: Mechanisms and Therapeutic Implications. Cells, 9(7). doi:10.3390/cells9071687

Pickrell, A. M., & Youle, R. J. (2015). The roles of PINK1, parkin, and mitochondrial fidelity in Parkinson’s disease. Neuron, 85(2), 257–273. doi:10.1016/j.neuron.2014.12.007

Poewe, W., Seppi, K., Tanner, C. M., Halliday, G. M., Brundin, P., Volkmann, J., … Lang, A. E. (2017). Parkinson disease. Nat Rev Dis Primers, 3, 17013. doi:10.1038/nrdp.2017.13

Qu, Y., Li, J., Qin, Q., Wang, D., Zhao, J., An, K., … Xue, Z. (2023). A systematic review and meta-analysis of inflammatory biomarkers in Parkinson’s disease. NPJ Parkinsons Dis, 9(1), 18. doi:10.1038/s41531-023-00449-5

Rahul, & Siddique, Y. H. (2021). Neurodegenerative Diseases and Flavonoids: Special Reference to Kaempferol. CNS Neurol Disord Drug Targets, 20(4), 327–342. doi:10.2174/1871527320666210129122033

Shaulian, E., & Karin, M. (2002). AP-1 as a regulator of cell life and death. Nat Cell Biol, 4(5), E131–136. doi:10.1038/ncb0502-e131

Simon, D. K., Tanner, C. M., & Brundin, P. (2020). Parkinson Disease Epidemiology, Pathology, Genetics, and Pathophysiology. Clin Geriatr Med, 36(1), 1–12. doi:10.1016/j.cger.2019.08.002

Singh, V. K., Sahoo, D., Agrahari, K., Khan, A., Mukhopadhyay, P., Chanda, D., & Yadav, N. P. (2023). Anti-inflammatory, anti-proliferative and anti-psoriatic potential of apigenin in RAW 264.7 cells, HaCaT cells and psoriasis like dermatitis in BALB/c mice. Life Sci, 328, 121909. doi:10.1016/j.lfs.2023.121909

Tolosa, E., Garrido, A., Scholz, S. W., & Poewe, W. (2021). Challenges in the diagnosis of Parkinson’s disease. Lancet Neurol, 20(5), 385–397. doi:10.1016/s1474-4422(21)00030-2

Vasudevan Sajini, D., Thaggikuppe Krishnamurthy, P., Chakkittukandiyil, A., & Mudavath, R. N. (2024). Orientin Modulates Nrf2-ARE, PI3K/Akt, JNK-ERK1/2, and TLR4/NF-kB Pathways to Produce Neuroprotective Benefits in Parkinson’s Disease. Neurochem Res, 49(6), 1577–1587. doi:10.1007/s11064-024-04099-8

Xiromerisiou, G., Hadjigeorgiou, G. M., Papadimitriou, A., Katsarogiannis, E., Gourbali, V., & Singleton, A. B. (2008). Association between AKT1 gene and Parkinson’s disease: a protective haplotype. Neurosci Lett, 436(2), 232–234. doi:10.1016/j.neulet.2008.03.026

Xu, L., Li, Y., Ji, J., Lai, Y., Chen, J., Ding, T., … Ge, W. (2022). The anti-inflammatory effects of Hedyotis diffusa Willd on SLE with STAT3 as a key target. J Ethnopharmacol, 298, 115597. doi:10.1016/j.jep.2022.115597

Ye, C., Zhang, B., Tang, Z., Zheng, C., Wang, Q., & Tong, X. (2024). Synergistic action of Hedyotis diffusa Willd and Andrographis paniculata in Nasopharyngeal Carcinoma: Downregulating AKT1 and upregulating VEGFA to curb tumorigenesis. Int Immunopharmacol, 132, 111866. doi:10.1016/j.intimp.2024.111866

Yu, Y., Zhou, M., Long, X., Yin, S., Hu, G., Yang, X., … Yu, R. (2023). Study on the mechanism of action of colchicine in the treatment of coronary artery disease based on network pharmacology and molecular docking technology. Front Pharmacol, 14, 1147360. doi:10.3389/fphar.2023.1147360

Zgorzynska, E., Dziedzic, B., & Walczewska, A. (2021). An Overview of the Nrf2/ARE Pathway and Its Role in Neurodegenerative Diseases. Int J Mol Sci, 22(17). doi:10.3390/ijms22179592

Zhang, W., Xiao, D., Mao, Q., & Xia, H. (2023). Role of neuroinflammation in neurodegeneration development. Signal Transduct Target Ther, 8(1), 267. doi:10.1038/s41392-023-01486-5

Zhao, L., Zhang, H., Li, N., Chen, J., Xu, H., Wang, Y., & Liang, Q. (2023). Network pharmacology, a promising approach to reveal the pharmacology mechanism of Chinese medicine formula. J Ethnopharmacol, 309, 116306. doi:10.1016/j.jep.2023.116306

Zheng, Y., Zhao, J., Chang, S., Zhuang, Z., Waimei, S., Li, X., … Zhao, G. (2023). β-Sitosterol Alleviates Neuropathic Pain by Affect Microglia Polarization through Inhibiting TLR4/NF-κ B Signaling Pathway. J Neuroimmune Pharmacol, 18(4), 690–703. doi:10.1007/s11481-023-10091-w

